# Cell-intrinsic regulation of presynaptic release site patterning

**DOI:** 10.64898/2026.10.02.756300

**Authors:** Jada D. Summerville, Elisa B. Frankel, Yongming Dong, Jihong Bai, Peri T. Kurshan

## Abstract

Functional neuronal connectivity depends not only on synaptic partner choice but also on the number, strength, and spatial distribution of release sites between partners. How these features are regulated remains poorly understood. Here, using the *C. elegans* DA9 motor neuron, we identify the immunoglobulin superfamily cell adhesion molecules SYG-1 and SYG-2 as cell-autonomous regulators of presynaptic release-site patterning. SYG-2 regulates presynapse number, size, and spacing, whereas SYG-1 primarily regulates spacing. Endogenous localization and cell-specific knockout analyses reveal that SYG-2 recruits SYG-1 to the presynaptic domain through previously unrecognized *cis* interactions. Endogenous structure-function studies further demonstrate that the SYG-2 intracellular domain is required for presynaptic organization and can promote presynapse formation independently of its extracellular domain. Genetic and targeted actin-disruption experiments implicate the adaptor protein NCK-1 and local actin remodeling in SYG-2 intracellular signaling. Together, these findings reveal how cell adhesion molecules can act cell-intrinsically to pattern presynaptic release sites after synaptic partners have been established.

## INTRODUCTION

The formation of functional neural circuits requires the precise organization of synaptic connections. Synaptogenesis involves multiple coordinated steps, including synaptic partner selection, pre- and postsynaptic assembly, and synapse maturation. Disruptions in any of these processes can lead to neurodevelopmental disorders (NDDs), underscoring the need for their tight regulation^1^. During development, structures such as the vertebrate retina^2,3^ and olfactory system^4^ form highly organized patterns of connectivity through selective synaptic partner choice. Consequently, much is known about mechanisms that determine which neurons connect with one another. Far less is known about how synapses are organized *between* established synaptic partners: whether the number, size and spatial distribution of individual release sites— features that shape synaptic strength and information transfer—are themselves regulated, and whether this organization is determined by cell-intrinsic mechanisms within the presynaptic neuron or through interactions with its postsynaptic partners.

*C. elegans* neurons form *en passant* synapses along their axons^5^, as do many neurons in the mammalian CNS^6^. The DA9 motor neuron forms approximately 25 dyadic synapses with body wall muscles and the inhibitory neuron VD13, with presynaptic release sites arranged in a highly stereotyped, regularly spaced pattern (**Figure 1A**). The boundaries of the DA9 presynaptic domain, and consequently the cells with which DA9 forms synapses, are established by extrinsic Wnt signals from epithelial cells in the tail^7,8^ and semaphorin/plexin signaling with the neighboring DA8 neuron^9^. In contrast, how individual presynaptic release sites are organized *within* this domain remains unknown. In particular, it is unclear whether their characteristic number, size and spacing are actively regulated and, if so, through what mechanisms.

**Figure 1:**
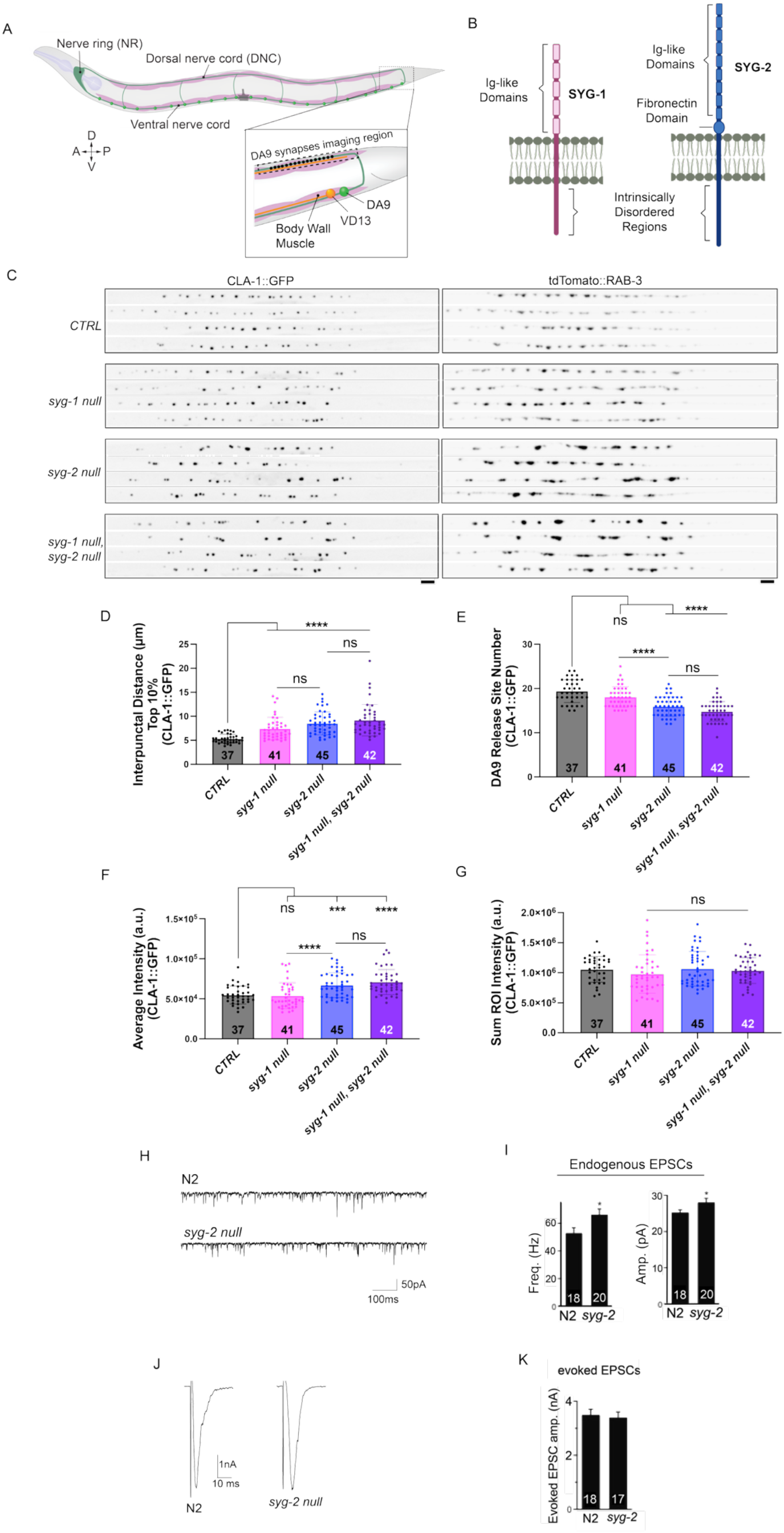
Loss of *syg-1* or *syg-2* disrupts presynaptic release site patterning in DA9. **A)** Schematic of *C. elegans* morphology. Inset highlights the location of DA9 presynapses. Dotted box within inset denotes the region shown in cropped images below and throughout. **B)** Schematic of SYG-1 and SYG-2 proteins. **C)** Straightened, cropped images of DA9 presynapes of four worms in CTRL, *syg-1 null*, *syg-2 null*, and *syg-1*, *syg-2* double mutants. Presynapses labeled with CLA-1::GFP and tdTomato::RAB-3 (*wyIs685* transgene). Scale bar = 4µm. **D)** Top 10% of interpunctal distances (measured as the 90^th^ percentile value of all interpunctal distances collected from each worm). **E)** Number of CLA-1::GFP puncta. **F)** Average CLA-1::GFP puncta intensity. **G)** Sum intensity of all the CLA-1::GFP puncta from each worm. **H)** Example traces of endogenous mEPSCs in wild type (N2) and *syg-2 null* animals recorded using patch clamp electrophysiology at the NMJ. **I)** Frequency and amplitude quantifications of endogenous mEPSCs. **J)** Example traces of evoked EPSCs in N2 and *syg-2 null* animals. **K)** Quantification of evoked EPSCs amplitude. Sample size (number of worms) indicated at the base of each bar. Each dot represents the value from a single worm. Error bars denote standard deviation.

In a forward genetic screen for regulators of presynaptic release site organization, we identified the evolutionary conserved immunoglobulin superfamily cell adhesion molecules (CAMs) SYG-1 and SYG-2. These CAMs were previously shown to mediate synaptic partner recognition in the *C. elegans* egg-laying circuit through *trans* interactions, with SYG-2 expressed by guidepost epithelial cells localizing presynaptic SYG-1 in adjacent neurons^10,11^. Here we identify a distinct, cell-autonomous role for SYG-1 and SYG-2 in organizing individual release sites within the DA9 presynaptic domain. Surprisingly, SYG-2 localizes presynaptic SYG-1 through extracellular *cis*, rather than *trans*, interactions. Moreover, the intracellular domain of SYG-2 is itself required cell-autonomously to regulate presynapse number, size, and spacing. Through extensive endogenous structure-function and cell-specific analyses we define the domains of SYG-1 and SYG-2 that are required for presynaptic release site organization. We further identify the SH2 adaptor protein NCK-1 and actin remodeling as mediators of this signaling pathway, reminiscent of signaling by the mammalian SYG-1/SYG-2 orthologs KIRREL and NEPRHIN in kidney cells^12^. Together, our findings reveal unexpected cell-intrinsic roles for SYG-1/SYG-2 *cis* interactions in patterning the stereotyped presynaptic release sites through local remodeling of the actin cytoskeleton. More broadly, they demonstrate that CAMs that specify synaptic partners in one context can be repurposed to organize individual release sites between established partners in another.

## RESULTS

### SYG-1 and SYG-2 are required for presynaptic release site patterning in DA9

To identify regulators of presynaptic release site patterning, we conducted a forward genetic screen in *C. elegans.* Presynaptic release sites were visualized by labeling active zone protein clarinet-1 (CLA-1::GFP), a functional homolog of mammalian piccolo^13^, along with RAB-3 (tdTomato::RAB-3), a synaptic vesicle associated protein^14^, specifically in the DA9 posterior motor neuron. In wild-type animals, DA9 forms around 25 *en passant* presynaptic release sites by the mid L4 larval stage of development. These release sites follow a stereotyped punctate pattern, with relatively uniform number, size and spacing (**Figure 1A, C**).

Whole genome sequencing and single nucleotide polymorphism^15^ mapping of one candidate with altered presynaptic patterning identified an early stop mutation in *syg-2*. To confirm that loss of *syg-2* was responsible for the phenotype, we used CRISPR to generate a full knock-out of the *syg-2* coding region, as well as that of its canonical binding partner, *syg-1* (**Figure 1B; Supplementary Figure 1A, B**). Analysis of the *syg-1* and *syg-2* null alleles using an automated synapse quantification tool developed in our lab^16^ revealed several quantifiable defects in presynaptic release site patterning. Because the more punctate CLA-1::GFP signal is easier to quantify, we used it for our quantitative analyses. Both mutants exhibited increased spacing between presynaptic release sites, characterized by larger gaps between CLA-1 puncta and quantified by comparing the top 10% of interpunctal distances compared to controls (*CTRL*; **Figure 1C-D**). Loss of *syg-2,* however, produced additional presynaptic patterning phenotypes, including a reduction in the number of presynaptic release sites (CLA-1 puncta) and, unexpectedly, an increase in the average intensity of the remaining CLA-1 puncta (**Figure 1C-F**). This increased intensity reflects an increase in CLA-1 molecules, interpreted as an increase in size of these diffraction limited structures. Despite this increase in average puncta intensity, summed puncta fluorescence (“sum ROI intensity”) was unchanged between *syg-2* mutants and controls (**Figure 1G**), indicating that presynaptic material is not lost but rather improperly redistributed among fewer sites. This phenotype is distinctive since most mutations that reduce synapse number also reduce synapse size. *syg-1, syg-2* double mutants were not significantly different from *syg-2* single mutants across these measures (**Figure 1C-G)**, consistent with *syg-1* and *syg-2* functioning in the same genetic pathway. However, alternative relationships cannot be excluded given the relatively mild *syg-1* phenotype compared to that of *syg-2*. This pronounced presynaptic patterning phenotype of *syg-2* mutants was evident throughout development, from L2 through adulthood, suggesting that SYG-2 is required to pattern the presynaptic domain while synapses are continuously added during larval growth (**Supplementary Figure 1C-D**).

To verify that the CLA-1::GFP phenotype in *syg-2* mutants was not an artifact of CLA-1 overexpression, we analyzed CLA-1 endogenously tagged with split-GFP^17^ (CLA-1::spGFP) in DA9. Similar to our exogenous integrated array marker, loss of *syg-2* resulted in a severe disruption of presynaptic patterning with reductions in endogenous CLA-1::spGFP puncta number and increased average intensity and interpunctal distances (**Supplementary Figure 1E-I**). These results validated the presynaptic patterning defects observed with the exogenous CLA-1::GFP marker.

To determine whether loss of *syg-1* and *syg-2* affected postsynaptic organization, we analyzed the muscle-specific nicotinic acetylcholine receptor subunit UNC-29^18^ endogenously tagged with RFP (UNC-29::RFP). Loss of either *syg-1* or *syg-2* had no detectable effect on the recruitment or distribution UNC-29::RFP (**Supplementary Figure 1J-L**). Collectively, these results suggest *syg-1* and *syg-2* preferentially affect presynaptic rather than postsynaptic organization.

To investigate the functional ramifications of the changes in presynaptic release site patterning in *syg-2* mutants, we performed patch-clamp electrophysiology recordings from the body wall muscles that receive synaptic input from these motor neurons. Despite having fewer presynaptic release sites, *syg-2* mutants exhibited significant increases in both frequency and amplitude of basal endogenous excitatory post-synaptic currents (EPSCs) (**Figure 1H-I**). This suggests that the remaining release sites in *syg-2* mutants have a higher probability of release, likely due to the increase in CLA-1 molecules at the active zone. The increase in endogenous EPSC amplitude indicated an increase in quantal size, which could arise from either presynaptic or postsynaptic mechanisms. Presynaptically this may reflect increased neurotransmitter content per vesicle, whereas postsynaptic possibilities include changes in receptor composition, function, localization, or alignment with release sites. Although we did not detect a change in postsynaptic receptor abundance, this alone does not distinguish among these possibilities (**Supplementary Figure 1J-L**). When neurons were stimulated to measure evoked release, we found that evoked EPSC amplitude was unchanged in *syg-2* mutants when compared to wild type (**Figure J-K**). Thus, despite a reduction in presynaptic release site number, overall evoked synaptic strength is maintained. Together with the increase in quantal size, these results suggest that compensatory changes in synaptic function help offset the decrease in presynaptic release sites. This aligns with the lack of any overt motor defects observed in *syg-2* mutants.

### SYG-2 functions cell-autonomously within DA9 to regulate presynaptic patterning

To determine from where SYG-1 and SYG-2 function to regulate DA9 presynaptic patterning, we first examined their expression by generating endogenous transcriptional reporters for each. Using CRISPR, we inserted a reporter cassette immediately upstream of the first exon of either *syg-1* or *syg-2*. The cassette included a GFP flanked by nuclear localization sequences (NLSs) and followed by a T2A ribosome-skipping sequence, allowing nuclear GFP and SYG-1 or SYG-2 to be translated from the same endogenous transcript (**Supplementary Figure 2A**). To identify specific cell types, we exogenously expressed cytosolic BFP under cell-specific promoters. Labeling DA9 with BFP revealed that both *syg-1* and *syg-2* were expressed in DA9 (**Supplementary Figure 2B**). Similarly, both *syg-1* and *syg-2* were also expressed in the postsynaptic body wall muscles (**Supplementary Figure 2C**). Interestingly, *syg-1*, but not *syg-2*, was expressed in a subset of inhibitory neurons including VD13, a postsynaptic partner of DA9 at its dyadic synapses (**Supplementary Figure 2D-F**). Taken together, these data suggest that SYG-1 and SYG-2 could function either cell-autonomously or non-cell-autonomously (**Figure 2A**).

**Figure 2:**
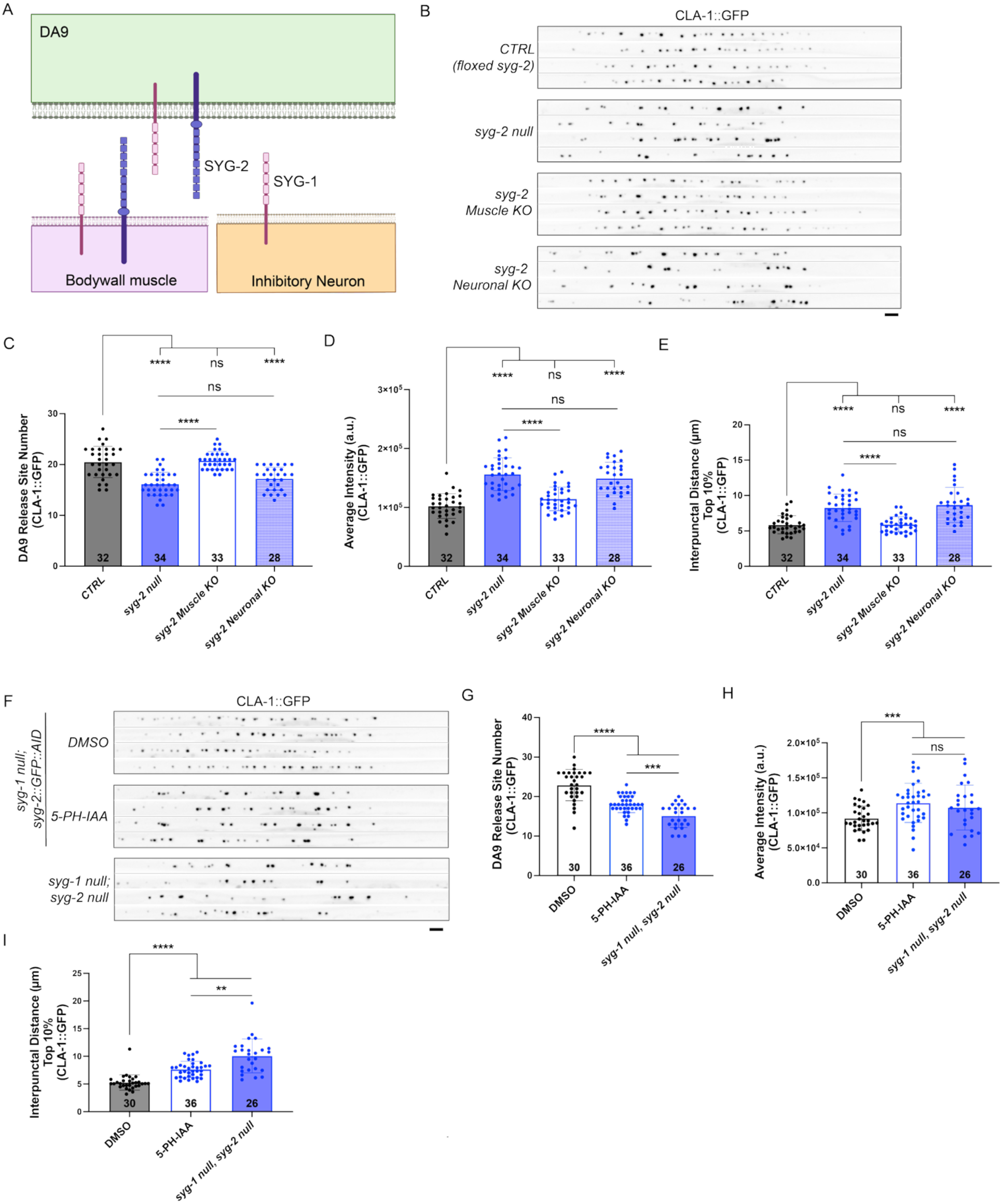
Loss of SYG-2 disrupts presynaptic patterning cell-autonomously. **A)** Schematic summary of SYG-1 and SYG-2 expression data. **B)** Straightened cropped images of CLA-1::GFP in DA9 of *CTRL, syg-2 null, muscle KO of syg-2 (pmyo-3::CRE),* and *DA9 neuron-specific KO of syg-2 (pmig-13::CRE)*. **C-E)** Quantification of CLA-1::GFP in *syg-2* conditional KOs. **F)** Images from L4 animals of CLA-1::GFP in DA9 from DMSO treated, 5-PH-IAA treated, and *syg-1 null, syg-2 null* mutant animals. **G-H)** Quantification of CLA-1::GFP from early neuronal degradation of SYG-2. Sample size (number of animals) indicated at base of bars. Scale bars = 4µm.

To determine whether SYG-2 functions presynaptically or postsynaptically to pattern DA9 presynaptic release sites, we performed cell-specific conditional knockouts using the Cre/LoxP system. Using CRISPR, we inserted loxP sites flanking the *syg-2* locus. Muscle-specific (i.e. postsynaptic) expression of Cre recombinase did not affect the patterning of DA9 presynaptic release sites compared to controls (**Figure 2B-E**), indicating that muscle-expressed SYG-2 is dispensable for DA9 presynaptic patterning. In contrast, DA9-specific (i.e. presynaptic) expression of Cre fully recapitulated the *syg-2 null* phenotype, resulting in fewer CLA-1 puncta, increased average puncta intensity, and increased interpunctal spacing (**Figure 2B-E**). Our results support a novel, cell-autonomous role for SYG-2 in DA9 presynaptic release site patterning.

Previous studies on synapse formation in the hermaphrodite-specific neuron (HSN) had demonstrated that SYG-2 expressed in non-neuronal guidepost epithelial cells delineates the location of HSN presynapses through *trans* interactions with its neuronal binding partner SYG-1^10,11^. We were therefore surprised to discover a neuronal cell-autonomous role for SYG-2 in DA9 synapse development. To confirm this result by assessing cell-autonomy in another, independent manner, we utilized the auxin-inducible degradation (AID) system^19^ to cell-specifically degrade SYG-2 in neurons. Using CRISPR, we endogenously tagged SYG-2 at its C-terminus with split GFP (7 copies of GFP11)^17^, with and without a 44 amino acid AID degron sequence. Neither the split-GFP alone nor the addition of the AID degron disrupted SYG-2 function in presynaptic patterning (**Supplementary Figure 3A-B**). To prevent any potential compensation by SYG-1 (and since the *syg-2* single mutants and *syg-1, syg-2* double mutants are indistinguishable), we performed the auxin-induced degradation in a *syg-1 null* background. We utilized a strain expressing TIR1 pan-neuronally together with an AID::BFP reporter allowing us to verify TIR1 activity. We independently confirmed SYG-2 degradation by crossing this strain into one expressing GFP1-10 in DA9 (**Supplementary Figure 3D**). To assess the effect of neuronal SYG-2 degradation, degron-tagged and control worms expressing CLA-1::GFP were grown on 50µM auxin analog (5-PH-IAA) or DMSO control plates. Neuronal degradation of SYG-2 during larval development recapitulated the major features of the *syg-2 null* phenotype (**Figure 2F-I**). The lack of full penetrance of some phenotypes was most likely due to the inability of auxin to penetrate the embryo, leading to incomplete abrogation of SYG-2 function at the earliest stages of development. Together with our conditional knockout experiments, these results provide independent evidence that SYG-2 functions in neurons to pattern DA9 presynaptic release sites.

### SYG-2 localizes SYG-1 to the presynaptic domain through *cis* interactions

Having established a neuron-specific role for SYG-2 in DA9 presynaptic release site patterning, we next examined its subcellular localization using our *syg-2* allele endogenously tagged with split GFP (SYG-2::spGFP). Imaging endogenous SYG-2 (spGFP) and CLA-1 (sp-mScarlet^17^) together in DA9 revealed that SYG-2 localizes throughout the synaptic domain, but is more clustered surrounding the active zone labeled with CLA-1 (**Figure 3A, A’**). We next examined the localization of SYG-1 within the DA9 neuron. We generated a *syg-1* allele endogenously tagged with split wrmScarlet (8 copies of wrmScarlet-11) at its C-terminus and visualized it together with CLA-1 endogenously tagged with spGFP^17^. SYG-1 also localized to the synaptic domain of DA9 but had a distinctly more punctate pattern of localization than SYG-2 (**Figure 3B, B’**). Like SYG-2, however, SYG-1 did not precisely colocalize with CLA-1 puncta but instead formed clusters around them. This distribution is a notable contrast to other CAMs we have studied in the past, such as neurexin, which clusters tightly with CLA-1 at active zones^20^. When directly compared in the same strain, SYG-1 and SYG-2 localizations largely overlapped (**Figure 3C, C’**), however, SYG-2 clusters frequently extended beyond the bounds of SYG-1 clusters. This indicates that although the two proteins colocalize within DA9’s synaptic domain, their subcellular distributions are partially distinct.

**Figure 3:**
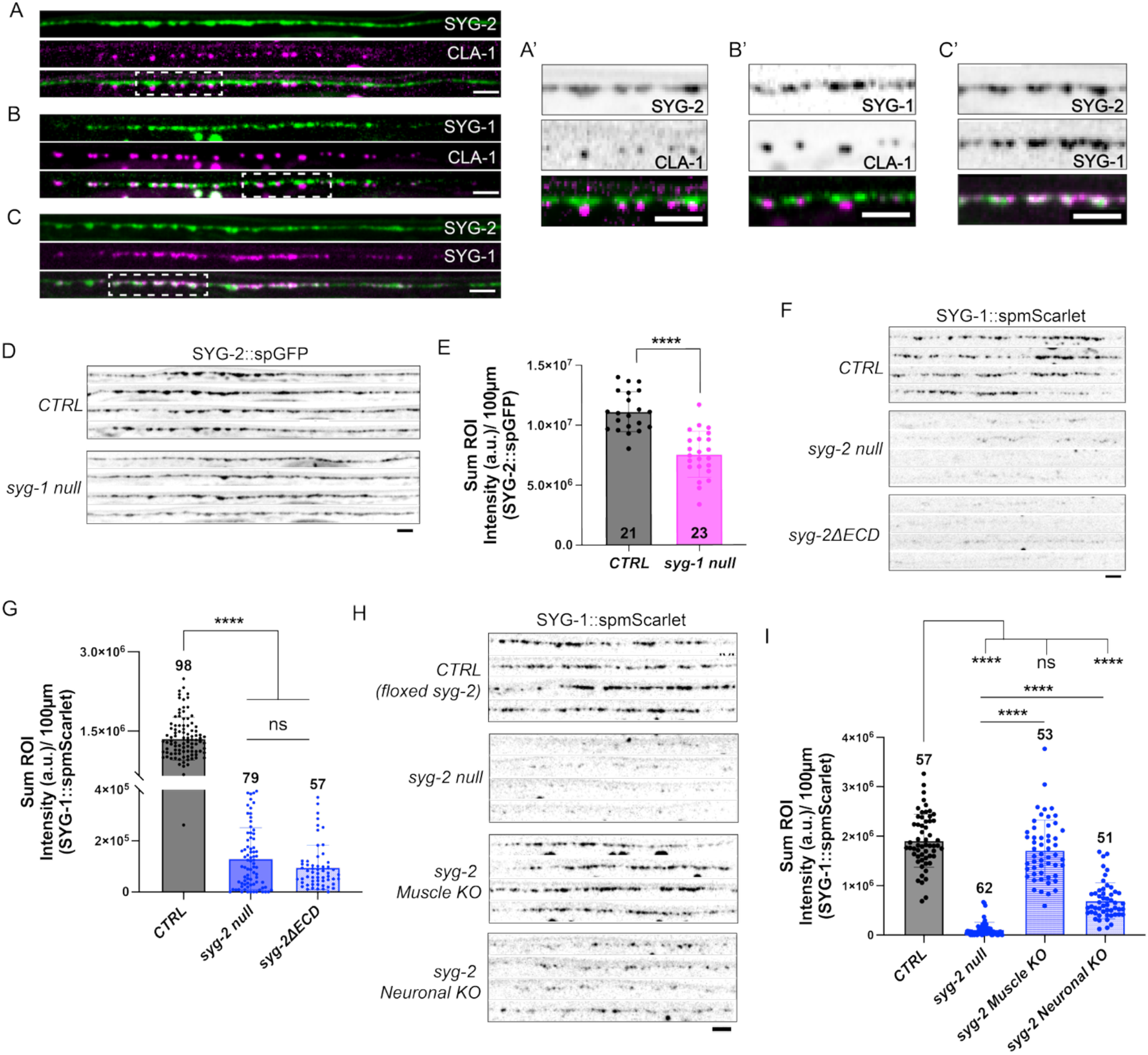
SYG-2 localizes SYG-1 at presynapses via *cis* interactions. **A)** Images of endogenous SYG-2::spGFP and CLA-1::spmScarlet, visualized cell-specifically in DA9 with an integrated dual split-FP transgene (*mizIs41* [*pmig-13::mScarlet1-10::SL2::GFP1-10*]^17^). **B)** Images of DA9 SYG-1::spmScarlet and CLA-1::spGFP. **C)** Images of DA9 SYG-2::spGFP and SYG-1::spmScarlet. **A’-C’)** Magnified regions from A-C (denoted by dotted boxes). **D)** Images of DA9 SYG-2::spGFP in *CTRL* and *syg-1 null* mutants. **E)** Quantification of SYG-2::spGFP sum intensity (normalized to image length). Sample size indicated at base of bars. **F)** Images of DA9 SYG-1::spmScarlet in *CTRL*, *syg-2 null*, and *SYGΔECD* mutants. **G)** Quantification of SYG-1::spmScarlet sum intensity (normalized to image length). **H)** Images of DA9 SYG-1::spmScarlet in *CTRL (floxed syg-2)*, *syg-2 null*, *syg-2* muscle knockout (pmyo-3::egl-13-NLS::wrmCRE) and *syg-2* neuronal knockout (pmig-13::egl-13-NLS::wrmCRE) animals. **I)** Quantification of SYG-1::spmScarlet sum intensity (normalized to image length) in H. Sample size indicated above error bars for G and I. Scale bars = 4µm.

Since our genetic analysis suggested that *syg-1* and *syg-2* function in the same pathway, and they are known direct binding partners^21^, we asked whether their localization in DA9 was dependent on one another. In *syg-1* mutants, SYG-2::spGFP intensity was reduced by approximately 40% (**Figure 3D-E**). However, the remaining SYG-2 retained its ability to localize and form clusters within the synaptic domain. This suggests that SYG-1 may be required to stabilize or maintain SYG-2 at the synapse but may not be required for SYG-2’s initial localization to the synaptic domain. In contrast, SYG-1 intensity was reduced by over 90% in *syg-2* mutants (**Figure 3F-G**), with approximately 10% percent of animals having no detectable fluorescence. Because both *syg-1* and *syg-2* are expressed in DA9, SYG-1 localization could potentially depend on either extracellular or intracellular interactions with SYG-2. To distinguish between these possibilities, we used CRISPR to delete the extracellular domain of SYG-2 (*syg-2ΔECD*) and analyzed SYG-1 localization. Loss of the SYG-2 extracellular domain resulted in a similar reduction of SYG-1 as the *syg-2* null (**Figure 3F-G**), demonstrating that SYG-2’s extracellular domain is required for SYG-1 localization. Taken together, these results reveal an asymmetric dependence between SYG-1 and SYG-2: SYG-2 can localize to the synaptic domain independently of SYG-1, whereas SYG-1 is almost completely dependent on extracellular interactions with SYG-2. These results also recontextualize how we interpret our original *syg-2 null* mutant, as loss of *syg-2* effectively disrupts SYG-1 localization, essentially creating a double mutant and potentially explaining why the *syg-2* null phenotype is much more severe than that of the *syg-1* null.

Since *syg-1* and *syg-2* are expressed in both the pre- and post-synaptic cells at the NMJ (DA9 and body wall muscle), they could interact either *in cis* or *in trans*. To distinguish between these possibilities, we utilized the FLP-FRT and Cre/LoxP conditional knockout systems to remove *syg-1* or *syg-2*, respectively, in a cell-specific manner and determine how loss of each protein affects the localization of the other. Using CRISPR, we inserted FRT sites flanking the *syg-1* locus and cell-specifically expressed a FLP recombinase from an extrachromosomal array. Loss of *syg-1* from the muscle resulted in an approximately 20% reduction in SYG-2 levels. Similarly, loss of *syg-1* from DA9 reduced SYG-2 by a similar magnitude (**Supplementary Figure 3G-H**). Neither cell-specific knockout fully recapitulated the *syg-1 null* phenotype, indicating that SYG-1 in DA9 and the muscle make comparable contributions to maintaining SYG-2 at the synaptic domain. These results suggest that SYG-1 can maintain SYG-2 localization through both *cis* and *trans* interactions. In contrast, using our floxed *syg-2* allele and cell-specific expression of Cre recombinase, we found that muscle knockout of *syg-2* had no detectable effect on SYG-1 localization (**Figure 3H-I**), while knockout of *syg-2* from DA9 resulted in a significant reduction in SYG-1 fluorescence similar to, although less severe than, that observed with complete loss of *syg-2* (**Figure 3H-I**). Taken together, these results reveal a novel *cis* interaction through which SYG-2 localizes SYG-1 to the DA9 presynaptic domain.

### SYG-2 can signal intracellularly in the absence of extracellular interactions

The presence of SYG-2 within the presynaptic domain, its cell-autonomous function in DA9 presynaptic patterning, and the more severe *syg-2 null* phenotype compared to that of *syg-1 null* mutants suggested that SYG-2 may signal through its intracellular domain to regulate presynaptic patterning. To test the requirements of the SYG-2 extracellular and intracellular domains, we generated endogenous domain deletions using CRISPR (**Supplementary Figure 1A**). We first deleted the extracellular domain (ECD) of SYG-2 (*syg-2ΔECD*), leaving only the transmembrane and intracellular domains (ICD) intact. Although *syg-2ΔECD* mutants exhibited defects in presynaptic patterning, these defects were not as severe as the *syg-2 null* (**Figure 4A-D**), indicating that the SYG-2 ICD retains partial function in the absence of the ECD. Compared with *syg-1* mutants*, syg-2ΔECD* mutants were more severe in measures of presynapse number and interpunctal distance, whereas average CLA-1::GFP intensity was intermediate between that of the *syg-1* and *syg-2 null* mutants. In contrast, deleting the SYG-2 ICD and replacing it with 7xGFP-11 (*syg-2ΔICD*), leaving the transmembrane domain and ECD intact, fully recapitulated the *syg-2 null* phenotype (**Figure 4A-D**). Together, these results demonstrate that the SYG-2 ICD is required for presynaptic patterning and can retain partial function independent of the ECD.

**Figure 4:**
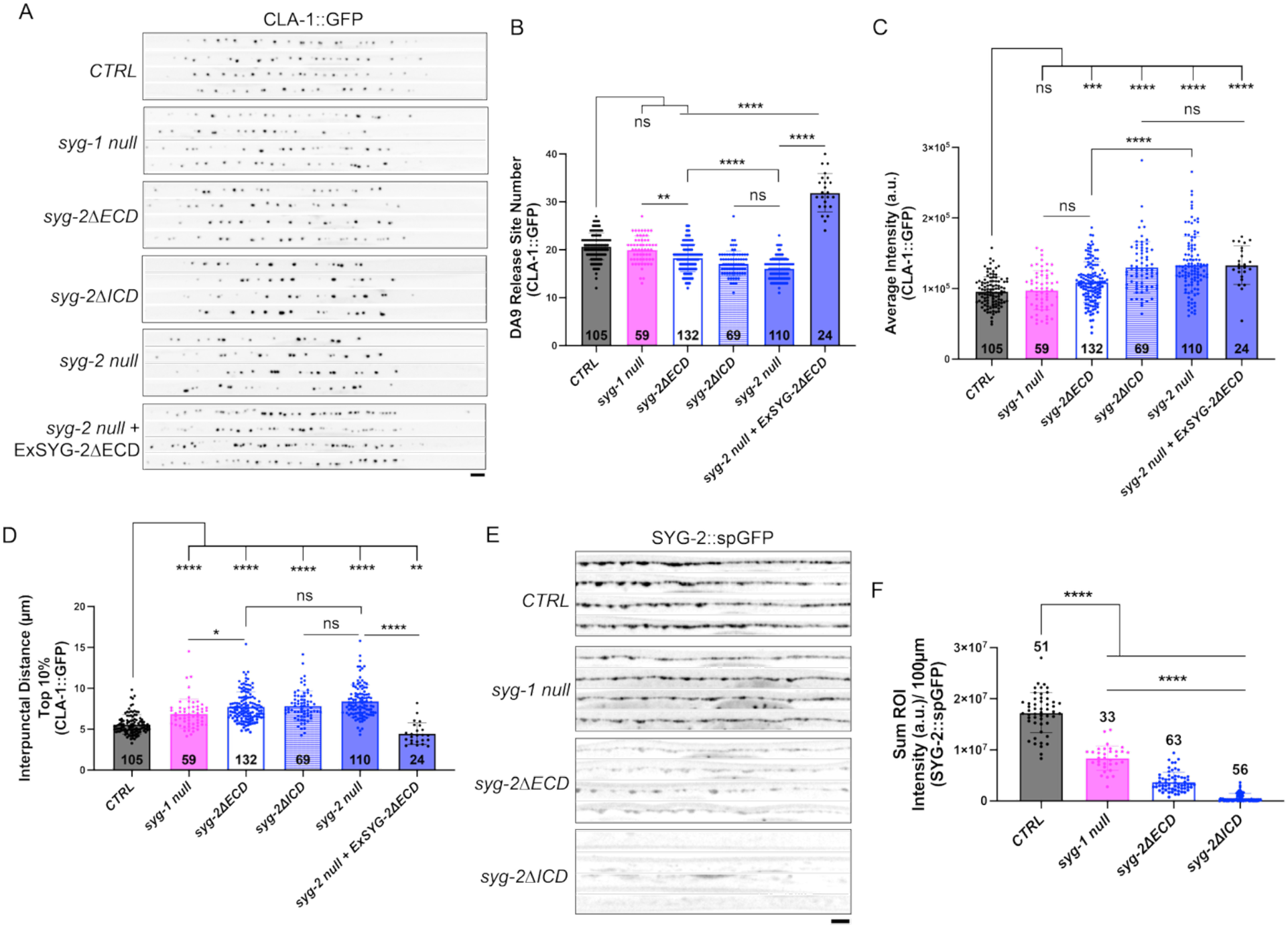
The SYG-2 intracellular domain can function independently of extracellular interactions and is required for SYG-2 localization. **A)** Images of CLA-1::GFP in DA9 of *CTRL, syg-1 null, syg-2ΔECD, syg-2ΔICD, syg-2 null* and *syg-2 null* with SYG-2ΔECD over-expressed extra-chromosomally in DA9 animals. B-D) Quantification of CLA-1::GFP. E) Images of DA9 specific SYG-2::spGFP in *CTRL*, *syg-1 null*, *syg-2ΔECD, and syg-2ΔICD* animals. F) Quantification of SYG-2::spGFP. Sample size (number of animals) indicated at base of bars. Scale bars = 4µm.

We next examined how these domains contribute to the localization of SYG-2 in DA9. To visualize endogenous SYG-2ΔECD localization, we introduced the ECD deletion into our split-GFP tagged SYG-2 strain. (SYG-2ΔICD already contained 7xGFP11 inserted at its C-terminus.) SYG-2ΔECDspGFP retained its ability to localize to the synaptic domain in DA9, although its fluorescence intensity was significantly reduced, to an even greater extent than that observed with the loss of *syg-1* (**Figure 4E-F**). SYG-2ΔICDspGFP, on the other hand, failed to localize to the synaptic domain (**Figure 4E-F**), demonstrating that the ICD is required for proper localization and/or trafficking of SYG-2 to the axon.

The partial function retained by *syg-2ΔECD* suggested that the SYG-2 ICD can regulate presynaptic patterning independently of extracellular interactions. To further test this, we overexpressed SYG-2 without its ECD in a *syg-2 null*. Strikingly, overexpression of SYG-2ΔECD resulted in an overabundance of CLA-1::GFP puncta (**Figure 4A-B**), providing further evidence that the SYG-2 ICD can promote presynaptic release site formation in the absence of its ECD. However, this overexpression did not rescue the increased CLA-1 puncta intensity phenotype of the *syg-2 null* mutants (**Figure 4C**), which suggests that the mechanisms that determine presynaptic release site number are separable from those that determine active zone size.

### SYG-1 and SYG-2 interact with n-WASP interacting protein, NCK-1

Our results suggesting that the SYG-2 ICD mediates cell-autonomous intracellular signaling prompted us to investigate potential downstream interactors. We used the Eukaryotic Linear Motif predictor^22^ to identify possible candidate motifs within the ICDs of SYG-1 and SYG-2 (data summarized in **Figure 5A**). Both ICDs contains a PDZ binding motif (PBM), which could interact with PDZ domain-containing active zone scaffold proteins^20^, Src-specific SH2-domain binding motifs (SH2 BMs) capable of interacting with SH2 domain-containing adaptor proteins, and a WIRS motif. The WIRS motif of SYG-1 had been implicated in regulating F-actin in the HSN neuron^23^. To test whether any of these motifs were required for SYG-2 function in presynaptic patterning, we used CRISPR to individually delete each motif and then quantified the effect on DA9 presynaptic release sites. Surprisingly, none of the individual motif deletions produced any defects (**Supplementary Figure 4A-B**). Simultaneous deletion of all three SYG-2 SH2 BMs also failed to illicit any defects in presynaptic patterning (**Supplementary Figure 4A-B**). Because SYG-1 shares many of the same motifs with SYG-2 and largely colocalizes with SYG-2 in the DA9 synaptic domain, we hypothesized that SYG-1 might act redundantly with SYG-2 and compensate for the loss of SYG-2 intracellular signaling. To test this, we combined the triple SYG-2 SH2 BMs deletion and a *syg-1 null* mutation. Loss of these motifs in conjunction with the loss of *syg-1* resulted in a reduced number of presynaptic release sites and an increase in interpunctal distance, resembling the *syg-2 null* phenotype (**Figure 5B-D**). Interestingly, however, the loss of the SH2 BMs did not produce an increase in the intensity of the remaining CLA-1::GFP puncta as observed in the *syg-2 null*, again supporting the notion that release site number and size may be mediated through different mechanisms (**Figure 5E**).

**Figure 5:**
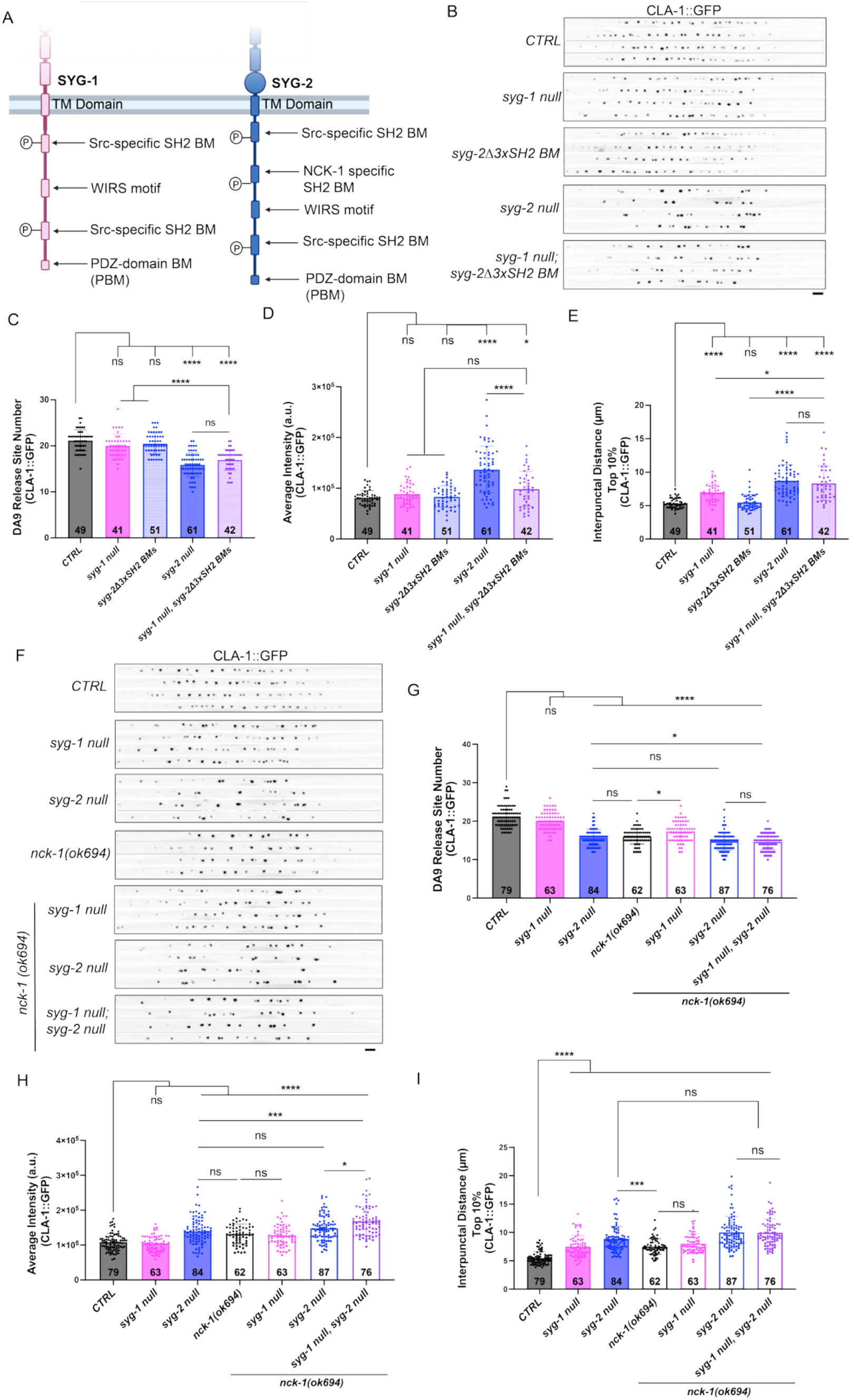
SYG-1 and SYG-2 interact with n-WASP interactor NCK-1. **A)** Schematic summary of motifs identified in SYG-1 and SYG-2 intracellular domains with the Eukaryotic Linear Motif predictor. **B)** Images of CLA-1::GFP in DA9 of *CTRL, syg-1 null, syg-2Δ3xSH2 BMs* (triple domain deletion), *syg-2 null*, and *syg-1 null, syg-2Δ3xSH2 BMs*. **C-E)** Quantification of CLA-1::GFP. **F)** Images of CLA-1::GFP in DA9 of *CTRL, syg-1 null, syg-2 null, nck-1 null*, and double and triple mutants. **G-I)** Quantification of CLA-1::GFP. Sample size (number of animals) indicated at base of bars. Scale bars = 4µm.

The mammalian orthologs of SYG-1 and SYG-2, KIRREL proteins and NEPHRIN, respectively, have been well characterized in organizing the slit diaphragm, a specialized cell junction in kidney podocytes^24–26^. NEPHRIN and KIRREL remodel the actin cytoskeleton by recruiting adaptor proteins, such as n-WASP interacting protein, NCK-1. The adaptor proteins bind to phosphorylated tyrosine sites on NEPHRIN/KIRREL. These interactions lead to the recruitment of the Arp2/3 complex to the membrane^12,27^. To investigate whether this pathway is conserved in presynaptic patterning for *C. elegans* SYG-1 and SYG-2, we examined DA9 presynaptic release sites in *nck-1*(*ok694*) mutants, a putative null allele consisting of a 1814 bp deletion^28^. Similar to *syg-2 null* mutants, *nck-1 null* mutants exhibited fewer CLA-1 puncta, increased puncta intensity, and increased interpunctal distance (**Figure 5F-I**). Furthermore, double mutants of *syg-2*; *nck-1* were not significantly different from *syg-2* single mutants in all aspects of presynaptic release site patterning. These non-additive phenotypes are consistent with SYG-2 and NCK-1 functioning in a shared genetic pathway.

Because of SYG-1’s ability to compensate for loss of SYG-2 intracellular signaling, we hypothesized that SYG-1 might also function through an NCK-1-dependent pathway. Indeed, the *syg-1*; *nck-1* double mutant was no more severe than the *nck-1* single mutant (**Figure 5F-I**). Similarly, the triple mutant *syg-1*, *syg-2; nck-1* did not differ significantly from the *syg-2*; *nck-1* double in release site number or interpunctal distance. Triple mutants, however, did exhibit a modest increase in average CLA-1::GFP intensity (**Figure 5F-I**). Overall, these non-additive genetic interactions are consistent with SYG-1 and SYG-2 functioning in an NCK-1-dependent pathway to regulate DA9 presynaptic release site patterning.

In HSN, SYG-1 has also been shown to remodel actin through its intracellular WIRS motif^23^, which sequesters F-actin binding scaffold protein NAB-1. NAB-1, in turn, was required for the localization of presynaptic active zone proteins SYD-1 and SYD-2^29^. To see whether NAB-1 functions similarly with SYG-2 in DA9, we examined DA9 presynaptic release sites in *nab-1* mutants. We generated a null allele of *nab-1* by deleting the entire coding region. Loss of *nab-1* resulted in defects in all aspects of presynaptic patterning including number, spacing, and intensity, but with some differences when compared to *syg-2 null* mutants (**Supplementary Figure 4C-F**). The *nab-1; syg-2 double mutant,* however, showed significantly greater defects than either of the single mutants in all parameters but CLA-1 intensity (**Supplementary Figure 4C-F**). These results suggest NAB-1 is important for presynaptic organization in DA9 but that it may function in a parallel pathway from SYG-2 for regulating presynapse number and spacing. In contrast, NAB-1 and SYG-2 may function in a shared pathway for regulating the amount of presynaptic material at individual release sites.

### Local disruption of actin polymerization around the SYG-2 ICD recapitulates the *syg-2 null* phenotype

The genetic interaction between *syg-2* and *nck-1* suggested that SYG-2 may regulate presynaptic patterning through local actin remodeling. DeActs are a class of peptides derived from *Salmonella enterica* which prevent or obstruct actin polymerization^30^ leading to the disassembly of actin filaments. To determine whether disruption of actin polymerization at SYG-2 specific sites could phenocopy loss of *syg-2*, we used CRISPR to fuse a DeAct peptide (SpvB) onto the C-terminus of our spGFP tagged SYG-2 allele. This allele failed to homozygose, potentially due to embryonic lethality arising from disruption of actin organization in muscle cells, which also express SYG-2. Indeed, using a lateral body wall muscle-specific GFP1-10 to visualize SYG-2::spGFP we observed that SYG-2 localized broadly throughout the muscles, and was present but not restricted to postsynaptic muscle arm termini (visualized with a membrane marker; **Supplementary Figure 5A**). To circumvent the lethality associated with a constitutive SYG-2::DeAct, we instead generated a FLP-dependent allele^31^ that enabled cell-specific expression of the DeAct-tagged protein (*syg-2::7xGFP11::FLPonDeAct*; hereafter *syg-2::FLPonDeAct*). In the absence of FLP-mediated recombination, this insertion did not disrupt SYG-2 function (**Supplementary Figure 5B-D**). We then introduced this allele into a *syg-1 null* background to prevent potential compensation from SYG-1. Activation of SYG-2::FLPonDeAct in DA9 resulted in a significant increase in both interpunctal distances and average intensity of CLA-1 puncta, recapitulating key features of the *syg-2 null* phenotype (**Figure 6A, C-D).** There was also a modest, but not statistically significant, reduction in CLA-1 puncta number (**Figure 6A-B**), perhaps due to the relatively late onset of expression of this DA9-specific promoter (approximately around L2/L3 stages). Together, these data support a model in which SYG-2 cell-intrinsically regulates presynaptic release site patterning through localized regulation of the actin cytoskeleton (**Figure 6E**).

**Figure 6:**
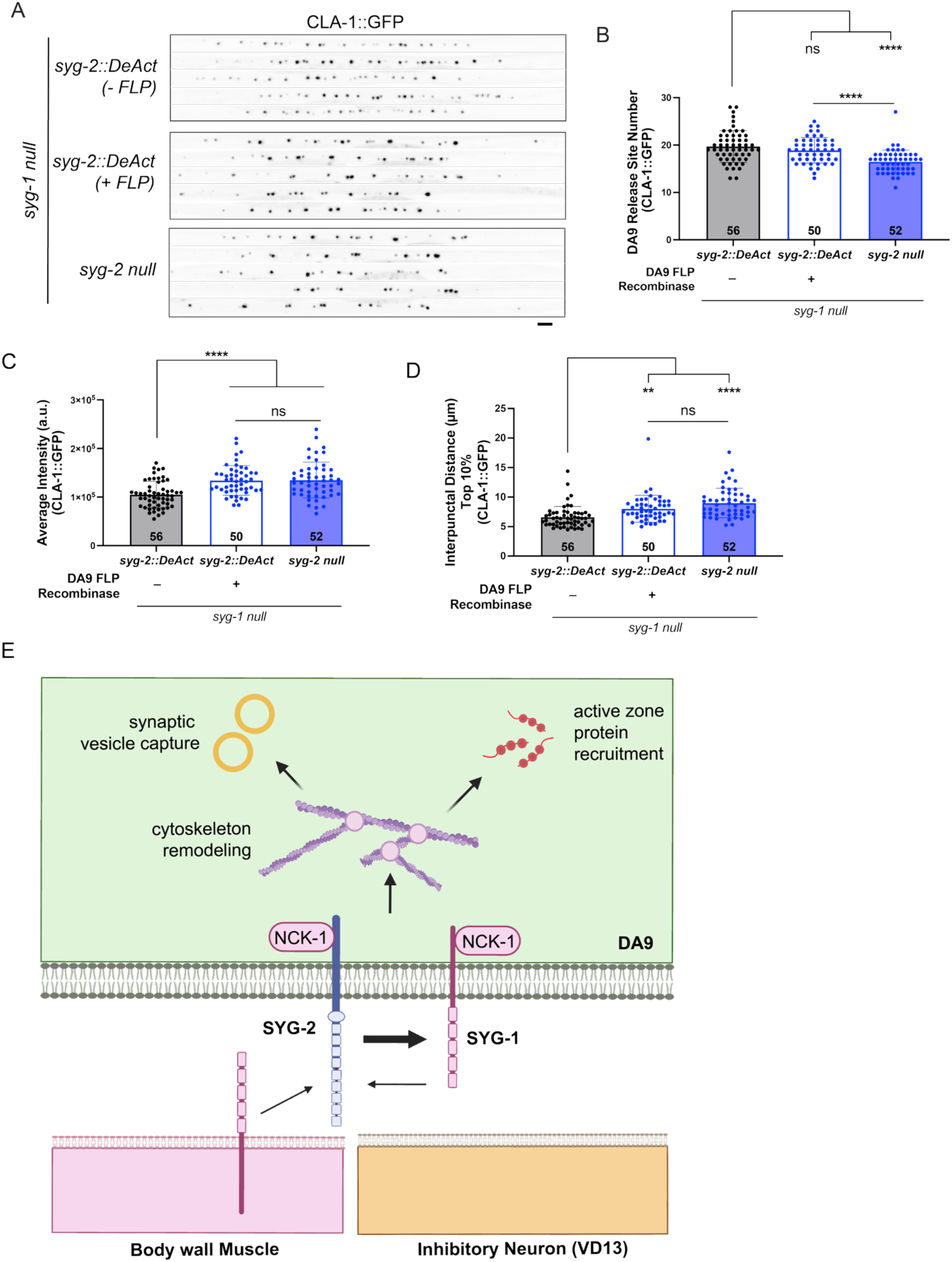
SYG-2 associated disruption of actin recapitulates the *syg-2* null phenotype. **A)** Images of CLA-1::GFP in DA9 of *syg-2::FLPonDeAct* (no FLP recombinase), *syg-2::FLPonDeAct* (DA9 specific FLP recombinase; *pitr-1::elg-13-NLS-FLP*), and *syg-2 null* animals. All strains are in a *syg-1 null* mutant background to prevent any possible compensation by SYG-1. **B-D)** Quantification of CLA-1::GFP. **E)** Model of SYG-1 and SYG-2 interactions, NCK-1 recruitment and local actin cytoskeleton remodeling in presynaptic release site patterning. Sample size (number of animals) indicated at base of bars. Scale bars = 4µm.

## DISCUSSION

Synaptic patterning has often been studied in the context of synaptic partner selection, as in the developing retina, or the subcellular targeting of synaptic inputs, as in cerebellar Purkinje cells^32^. However, whether cell-intrinsic mechanisms regulate the distribution of presynaptic release sites between established synaptic partners remains poorly understood. Here we identify a cell-autonomous role for the highly conserved cell adhesion molecule SYG-2 in patterning the number, size, strength, and spacing of presynaptic release sites in the DA9 motor neuron, with functional ramifications for basal release properties. SYG-2’s binding partner, SYG-1, is also required for proper presynaptic spacing, but not for presynapse number or size. SYG-1 and SYG-2 are mutually but asymmetrically dependent on one another for their localization at the presynaptic domain: SYG-1 localization is strongly dependent on SYG-2, whereas SYG-2 localization is only partially dependent on SYG-1. Notably, SYG-2 localizes SYG-1 through extracellular *cis*, rather than *trans*, interactions, revealing an unexpected mode of action for these cell adhesion molecules. We also demonstrate that SYG-2 can signal through its intracellular domain independently of its extracellular domain. Our genetic analyses place SYG-2 and the SH2 adaptor protein NCK-1 in a common pathway, while targeted disruption of actin polymerization at the SYG-2 intracellular domain recapitulates key features of the *syg-2 null* phenotype. Together, these findings support a model in which cell-autonomous SYG-1/SYG-2 signaling organizes presynaptic release sites through local remodeling of the actin cytoskeleton (**Figure 6E**).

The precise mechanism by which SYG-2-mediated actin remodeling serves to pattern presynaptic release sites remains unclear. One possibility is that SYG-2 promotes the formation of actin-rich ‘landing sites’ that capture presynaptic components transported from the soma along microtubules by kinesin motors^33–35^. Alternatively, SYG-2 might regulate the redistribution of presynaptic material within the synaptic domain through actin-dependent inter-synaptic vesicle exchange^36,37^, which allows presynaptic material to be shared between synaptic boutons^38^. In this model, SYG-2-dependent actin ‘bridges’ could facilitate trafficking of presynaptic material between neighboring boutons, thereby limiting presynapse size. Consistent with this possibility, SYG-2 is distributed throughout the synaptic domain, including between presynaptic sites. However, overexpression of SYG-2ΔECD demonstrates that ectopically increasing the number of presynaptic CLA-1 puncta does not result in elimination of the abnormally large puncta in *syg-2 null* mutants (**Figure 4**), thus arguing against a simple model in which presynaptic release site number and size are coupled. Thus, the increased accumulation of CLA-1 at individual release sites in *syg-2* mutants is unlikely to arise solely as a compensation for reduced presynapse number and instead suggests that SYG-2 may independently regulate presynapse number and size.

SYG-1 has been previously shown to delineate the synaptic domain in the *C. elegans* HSN neuron through actin remodeling^23^. In HSN, SYG-1-dependent F-actin remodeling was shown to recruit the adaptor protein NAB-1, which in turn promotes localization of the active zone scaffolds SYD-1 and SYD-2^29^. In DA9, loss of *nab-1* also disrupts presynaptic patterning, but genetic interactions between *nab-1* and *syg-2* suggest that they function in distinct pathways to regulate presynapse number and spacing. Thus, although actin remodeling appears to be a shared feature of SYG-1/2-dependent synapse organization, the downstream mechanisms may differ between neuronal contexts.

The cell-autonomous role of SYG-2 in DA9 contrasts with its previously described role in patterning HSN presynapses. In HSN, SYG-2 functions non-cell-autonomously from guidepost epithelial cells, where it interacts *in trans* with SYG-1 localized on the HSN axon to specify the location of synapse formation^10,11^. In DA9, in contrast, neuronal SYG-2 localizes SYG-1 through extracellular *cis* interactions and signals through its own intracellular domain. Moreover, the SYG-2 intracellular domain retains activity in the absence of its extracellular domain, indicating that extracellular interactions are not strictly required for its signaling function. This *cis* interaction between SYG-1 and SYG-2 reveals an unexpected mode of action for proteins previously characterized primarily as *trans-*cellular adhesion molecules. Together these findings illustrate how the same pair of cell adhesion molecules can employ distinct interaction modes and signaling mechanisms to regulate different aspects of synapse development.

Cell adhesion molecules regulate multiple aspects of synaptogenesis including partner recognition, pre- and postsynaptic assembly, and synapse maturation^39^ and disruption of these proteins is associated with numerous neurodevelopmental disorders^40,41^. CAMs can organize synapses both by directly recruiting synaptic components^20,42,43^ and indirectly by remodeling the actin cytoskeleton^23,44^. SYG-1 and SYG-2 are highly conserved across both invertebrates and mammals in both neuronal and non-neuronal contexts. In *Drosophila,* SYG-1 and SYG-2 orthologs Kirre and Sticks and Stones (Sns) mediate diverse processes including myoblast fusion^45^, neural wiring of the neuromuscular^46^ and the olfactory systems^47^, and nephrocyte development^48,49^. At the Drosophila NMJ, SYG-2 ortholog Sns regulates NMJ morphology through interactions with a Type IIa receptor-like protein tyrosine phosphatase, rather than the SYG-1 ortholog^46^. Mammalian KIRREL1-3 and NEPHRIN, orthologs of SYG-1 and SYG-2, respectively, are best characterized for their roles in formation of the kidney slit diaphragm^25,26,50^, but KIRREL proteins also have important functions in the nervous system. KIRREL2 and KIRREL3 mediate attractive homophilic and repulsive heterophilic interactions that contribute to axon targeting in the mouse olfactory system^51–53^, and KIRREL3 also regulates connectivity within olfactory glomeruli^54^. KIRREL3, which also regulates synaptic patterning at mossy fiber synapses of the hippocampus through homophilic binding^55^, has been associated with autism and intellectual disability^56^, and some NDD-associated variants reside within its intracellular domain^57^, underscoring the importance of understanding KIRREL-family intracellular signaling. Our findings identify a novel cell-autonomous SYG-2 signaling pathway that organizes individual presynaptic release sites and reveal that proteins best known for specifying cell-cell interactions can also act intrinsically within a neuron to pattern synaptic connectivity after partner selection has occurred.

## MATERIALS AND METHODS

### *C. elegans* maintenance

All C. elegans were maintained at 20-23°C on nematode growth medium (NGM) plates seeded with OP50 E. coli as standard protocols^58^. Transgenic strains carrying extrachromosomal arrays were generated by injecting animals with 1-20ng/µL of plasmid alongside co-injection markers podr-1::RFP (100-200ng/µL), pmyo-2::mCherry (1.5-2ng/µL) or *rol-6* (50ng/µL). All strains used in this study are listed in Supplementary Table 1.

### Molecular biology

All plasmids were constructed using Gibson isothermal assembly^59^ and cloned into ΔpSM vector backbone.

### Generation of CRISPR-Cas9 edited strains

Endogenous gene edits of *syg-1* and *syg-2* were created by gonadal injection of CRISPR-Cas9 protein complexes. CRISPR reagents were ordered from IDT (Cas9, tracrRNA, ssDNA repair templates, and crRNA). Insertions larger than 40bp were either ordered as G-block fragments from IDT or cloned from existing plasmids. Repair templates for these larger insertions were synthesized with 150-200 bp homology arms cloned from worm DNA and assembled with the insertions using Gibson Assembly. Smaller edits (<40 bp) were ordered from IDT as 4nmol ultramers (200 bp) and designed with homology arms of 80-100 bp. We utilized a co-CRISPR strategy^60^, targeting *dpy-10*. dsDNA from PCR products for larger insertions were injected at a final concentration of 100ng/µL while ssDNA repair templates (4nmol ultramer) were injected for a final concentration of 2.5 µM. CRISPR edits were genotyped through PCR and confirmed by sequencing and outcrossed.

### Confocal Microscopy

Images of L4 animals were collected at room temperature, staged within a 2 hour developmental window imaged at L4 stages L4.4-L4.6^61^ unless indicated otherwise. Images of L1 animals were collected ∼9 hours after eggs were laid. L2 animals were collected ∼27 hours after eggs laid. L3 animals were collected after ∼35 hours after eggs laid. Young adult animals were singled at L4 stage and placed at 20°C overnight and collected the following day. 20-40 hermaphrodites were mounted onto glass slides layered with 10% agarose pads and paralyzed with 20mM Levamisole (Sigma Aldrich) dissolved in M9 buffer. Slides were sealed with an 18X18mm cover slip and Vaseline. Animals were imaged with Zeiss Plan-Apochromat 63x/1.4NA oil immersion objective. Images were acquired on a spinning disk confocal microscope (3i) equipped with an Evolve 512 EMCCD camera (Photometrics) and a CSU-X1 M1 spinning disk (Yokogawa) mounted on an inverted Zeiss Axio Observer Z1 microscope. Confocal z-stacks of the DA9 axon or dorsal nerve cord were collected with 0.27µm z steps over a 6 µm range.

### Confocal Image Processing

Confocal images were processed in Fiji. Cropped straightened images of the DA9 synaptic domain were aligned from the turn of the commissure. Images of SYG-1::spmScarlet and SYG-2::spGFP were cropped from the start of detectable fluorescence to the last detectable cluster/puncta. Images of endogenous CLA-1::mScarlet were traced from the posterior end of detectable dorsal nerve cord puncta to the end of the captured image. Presynaptic release sites and other fluorescently labeled proteins were quantified using the MATLAB application developed in our lab, WormSNAP^16^.

### Auxin Degradation Experiments

Auxin plates were generated from a 1mM concentrated solution of auxin analog, 5-PH-IAA. 5-PH-IAA was first resuspended in DMSO to create a 50mM stock solution that was then diluted to 1mM with M9 buffer. 1mM solution of 5-PH-IAA was added to 12 mL NGM plates for a final concentration of 50µM 5-PH-IAA. Plates were left to dry overnight under foil and then used immediately or stored at 4°C for up to one month. For control plates, DMSO of equivalent concentration was added to NGM plates. Eight adult animals were placed on auxin and control plates and allowed to lay eggs for 2-3 hours. Adults were removed and the plates were placed at 20°C for three days prior to imaging. Animals were imaged at L4 stage (L4.4-L4.6).

### *C. elegans* Electrophysiology

Electrophysiological recordings were performed on day 1 adult hermaphrodite worms as described previously^62^. Animals were immobilized on Sylgard-coated coverslips with tissue adhesive (Histoacryl Blue, Braun) and dissected in extracellular solution by making a dorsolateral incision with a sharpened tungsten needle. The gonad and intestine were removed by gentle suction through a glass pipette. To expose the ventral nerve cord and adjacent body wall muscle, the cuticle flap was reflected and secured with glue. Preparations were then transferred to a fixed-stage upright microscope (BX51WI, Olympus) equipped with a 60x water-immersion objective. The integrity of the anterior ventral body wall muscles and ventral nerve cord was verified by differential interference contrast microscopy.

### Quantification and Statistical Analysis

Graphs were constructed using Prism (v11.0.2, GraphPad Software Inc.). All quantification is presented with mean and error bars representing standard deviation. Sample size (number of animals) is denoted at the base of bars or above error bars. After collecting all interpunctal distances, we used the 90^th^ percentile value of each worm to determine the top 10%. Datasets from multiple imaging sessions with similar controls and the same WormSNAP^16^ parameters were combined. Data was analyzed in Prism. Unpaired t-tests with Welch’s correction were performed for single pairwise comparisons and Brown-Forsythe and Welch ANOVA tests with Dunnett’s T3 multiple comparisons test was performed when comparing multiple groups. ns = not significant; ****p<0.0001; *** p<0.001; ** p<0.01; * p<0.05.

## Acknowledgments

We would like to thank members of the Kurshan lab as well as Hannes Buelow, Scott Emmons and David Hall for helpful feedback and discussion. We thank Kota Mizumoto and Shaul Yogev for sharing strains. This work was funded by NINDS F31NS134338 (JDS) and NINDS R01NS123645 (PTK).

---

**Supplementary Figure S1:**
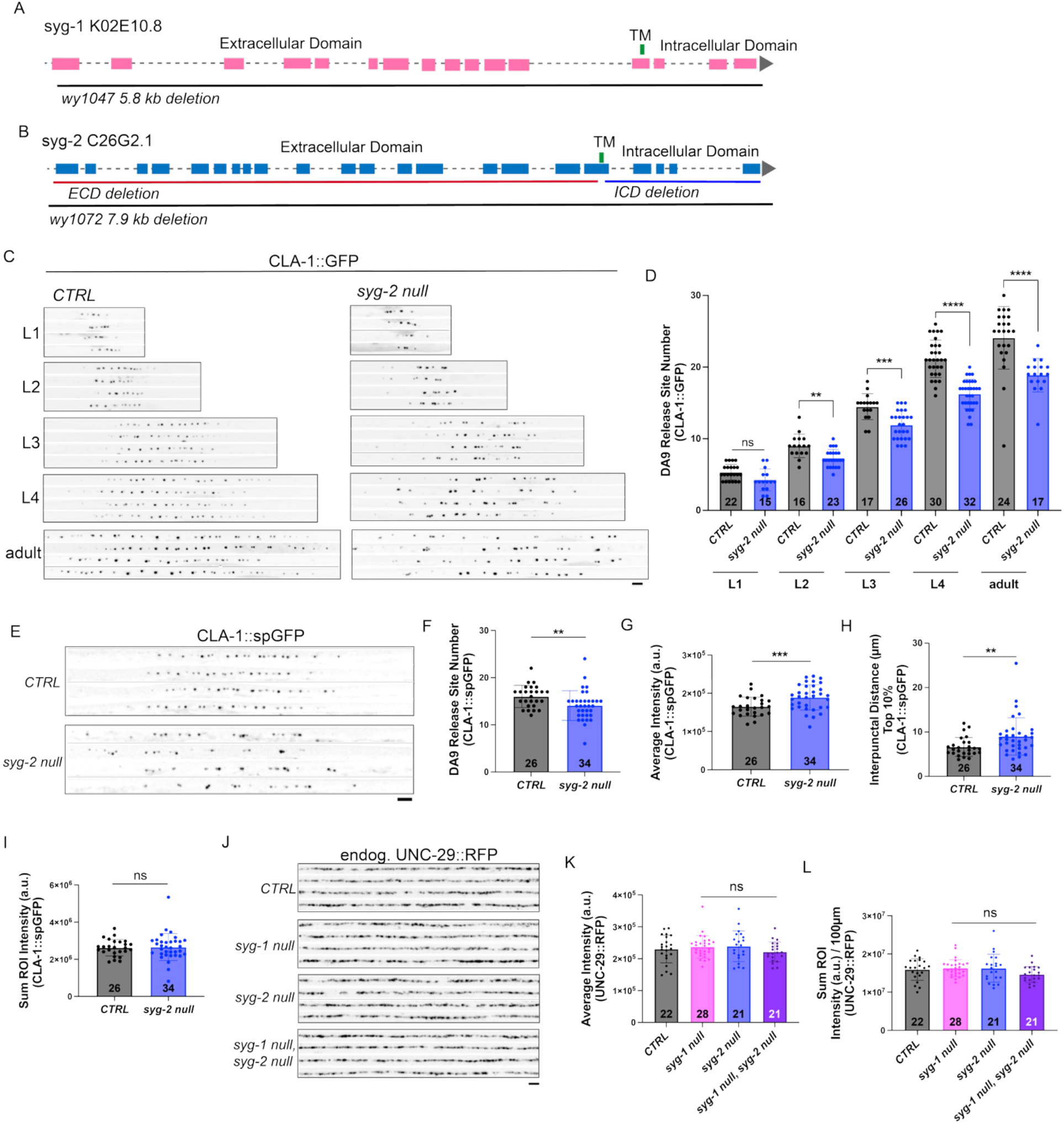
*syg-2* DA9 patterning defect is apparent with endogenously tagged active zones, throughout development, and is specific to presynapses. **A-B)** Schematics of *syg-1* and *syg-2* genetic locus, protein domains and deletion alleles. Schematics adapted from Wormbase sequences. **C)** Images of CLA-1::GFP (*wyIs685*) in *CTRL* and *syg-2 null* animals throughout development (L1-adult). **D)** Quantification of CLA-1::GFP, *CTRL*s compared to mutants of the same age. **E)** Images of endogenous split-GFP tagged CLA-1 in DA9 (CLA-1::spGFP) in *CTRL* and *syg-2 null* animals. **F-I)** Quantification of endogenous CLA-1::spGFP from C. **J)** Images of dorsal muscles with endogenous UNC-29::RFP labeling AChRs. **K-L)** Quantification of UNC-29::RFP. Sample size (number of animals) indicated at base of. Scale bars = 4µm.

**Supplementary Figure S2:**
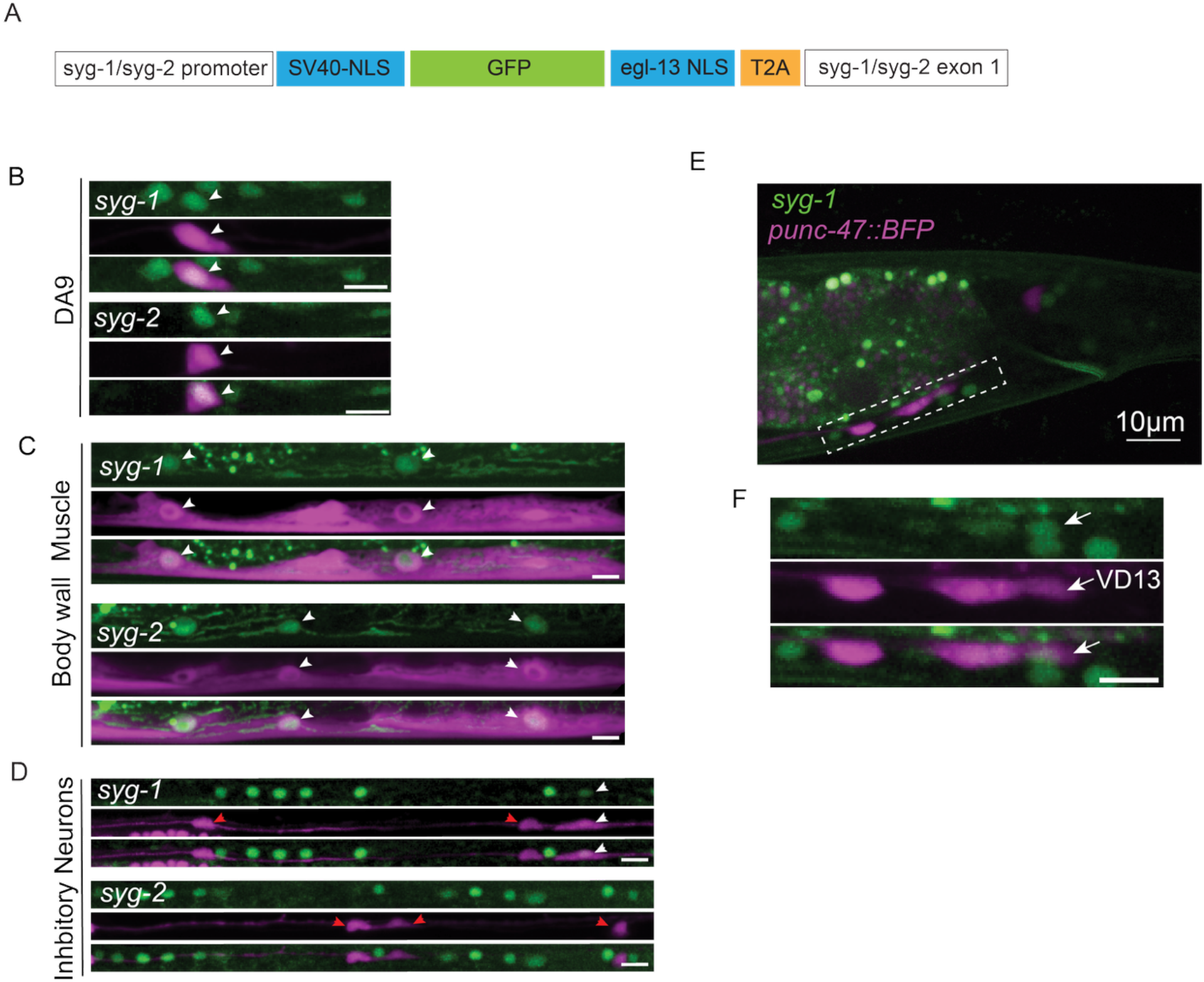
*syg-1* and *syg-2* are expressed in DA9 and its postsynaptic partners. **A)** Schematic of transcriptional reporter CRISPR insertion. **B)** Images of *syg-1* and *syg-2* transcriptional reporters co-localizing with DA9 (*pmig-13::elg-13-NLS-BFP*). Arrowheads denote DA9 nucleus. **C)** Images of *syg-1* and *syg-2* transcriptional reporters co-localizing with body wall muscle cells (*pmyo-3::elg-13-NLS-BFP*). Arrowheads denote muscle nuclei. **D)** Images of *syg-1* and *syg-2* transcriptional reporters co-localizing with inhibitory neurons (*punc-47::elg-13-NLS-BFP*). White arrowheads denote inhibitory neuron nuclei co-localizing with the *syg-1* transcriptional reporter, red arrowheads denote inhibitory neuron nuclei with no co-localization. **E)** Tail region of an animal with *syg-1* transcriptional reporter and inhibitory neurons labeled. **F)** Magnification of dotted box in E. Arrow denotes the most posterior inhibitory neuron, VD13. Scale bars = 5µm unless indicated otherwise. BFP fluorescence shown as magenta.

**Supplementary Figure S3:**
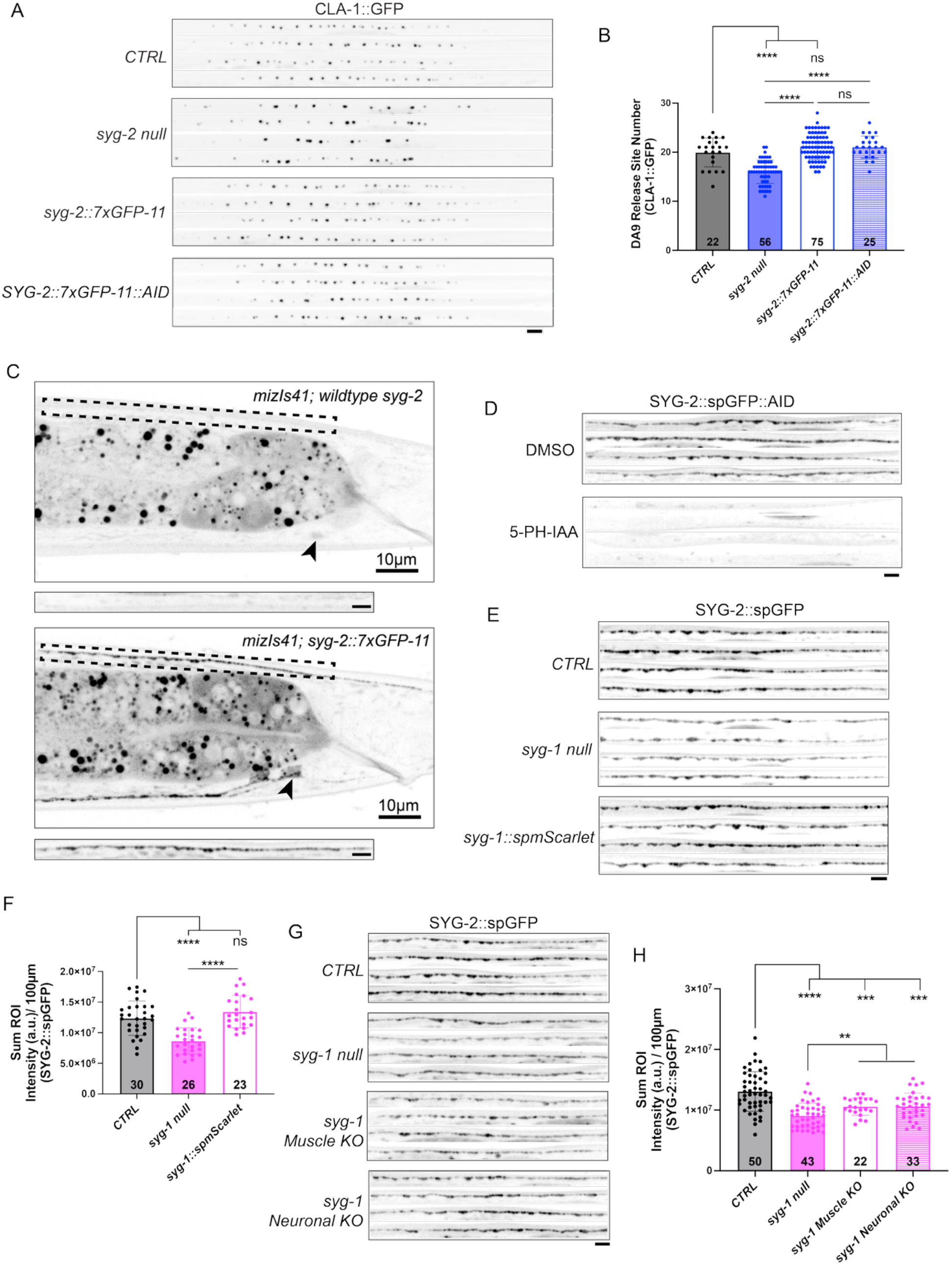
Endogenously tagging SYG-1 and SYG-2 with split-fluorophore and degron tags does not impact their function. **A)** Images of DA9-specific CLA-1::GFP (*wyIs685*) in *CTRL*, *syg-2 null*, *syg-2::7xGFP11* with no GFP1-10 expression, and *syg-2::7xGFP11::AID* (no auxin) animals. **B)** Quantification of CLA-1::GFP from C. **C)** Tail images of animals with DA9-specific GFP1-10 (*mizIs41*) and either wild type *syg-2* or *syg-2::7xGFP11*. Dashed box shows presynaptic domain shown in inset. Arrowhead indicates DA9 cell body. **D)** Images of DA9-specific SYG-2::spGFP::AID in DMSO and auxin (5-PH-IAA) treated SYG-2::spGFP::AID animals. **E)** Images of DA9 specific SYG-2::spGFP in *CTRL*, *syg-1 null* and *syg-1::spmScarlet* animals. **F)** Quantification of SYG-2::spGFP. **G)** Straightened cropped images of SYG-2::spGFP in *syg-1 null*, and *syg-1* muscle knockout (*pmyo-3::egl-13-NLS-FLP*) and *syg-1* DA9 neuronal knockout (*pitr-1::egl-13-NLS-FLP*) animals. **H)** Quantification of SYG-2::spGFP in I. Sample size (number of animals) indicated at base of bars. Scale bars = 4µm unless indicated otherwise.

**Supplementary Figure S4:**
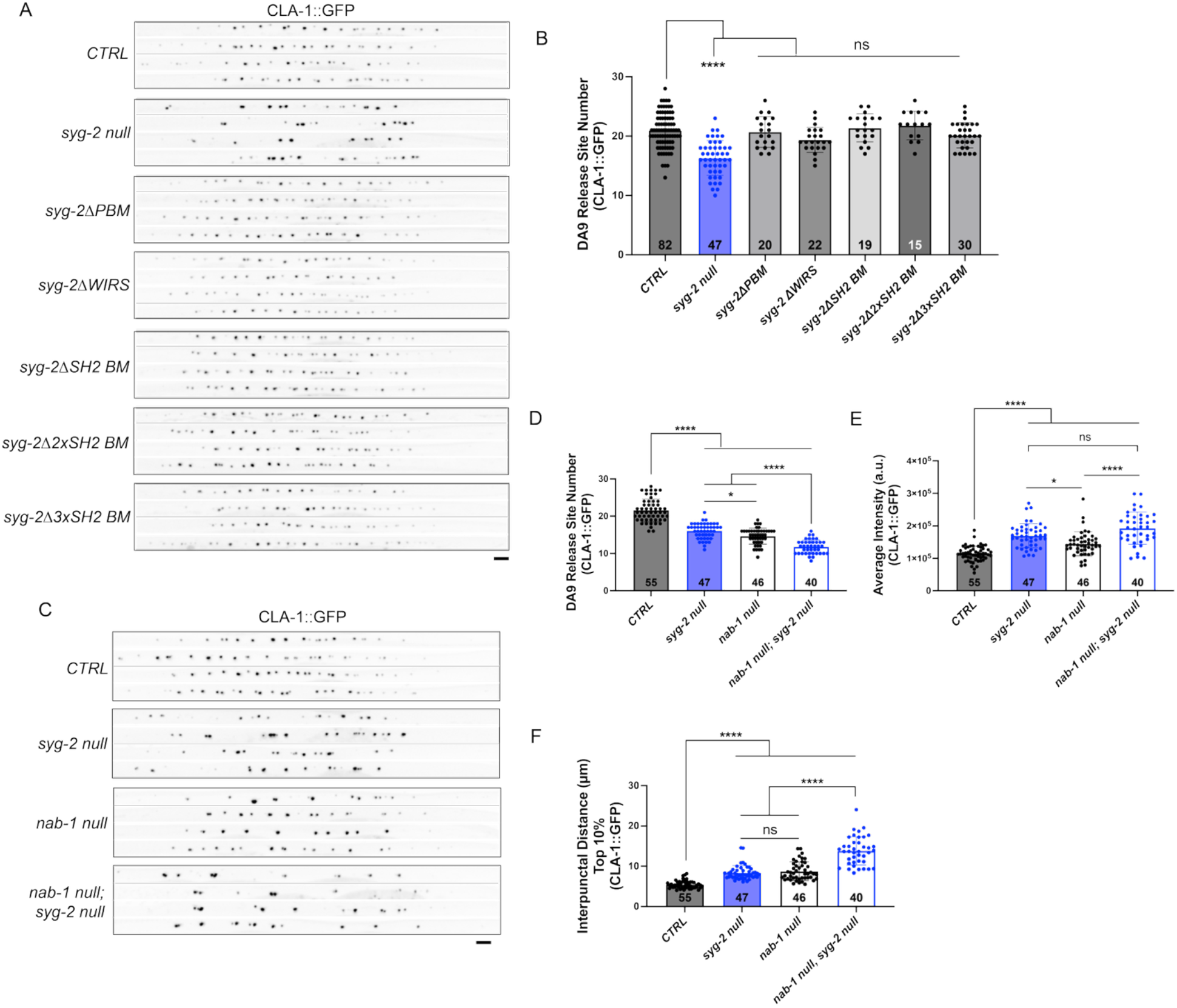
SYG-2 intracellular interactions. **A)** Images of DA9 specific CLA-1::GFP in *CTRL*, *syg-2 null*, and *syg-2* sub-domain mutants. **B)** Quantification of CLA-1::GFP. **C)** Images of DA9 specific CLA-1::GFP in *syg-2* and *nab-1* mutants. **D-F)** Quantification of CLA-1::GFP from C. Sample size (number of animals) indicated at base of bars. Scale bars = 4µm.

**Supplementary Figure S5:**
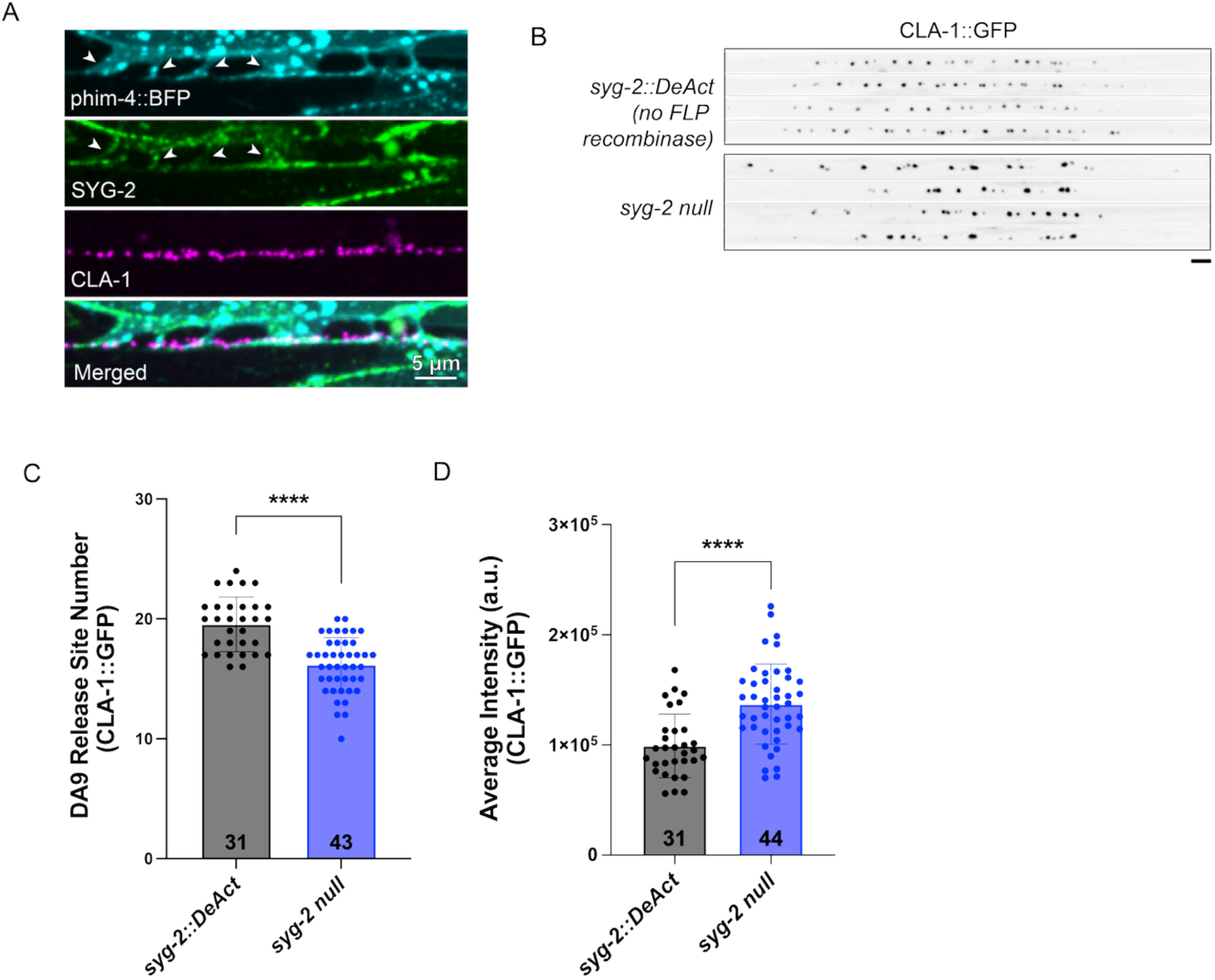
SYG-2 DeAct tag validation. **A)** Images of muscle arms (*phim-4::BFP-PH*), endogenous SYG-2::spGFP in the muscle (*phim-4::GFP1-10*), and endogenous CLA-1::spmScarlet. Arrowheads denote body wall muscle arms contacting neurons in the dorsal nerve cord. **B)** Images of DA9-specific CLA-1::GFP in *syg-2::FLPonDeAct* (no FLP recombinase expression) and *syg-2 null*. **C-D)** Quantification of CLA-1::GFP from B. Sample size (number of animals) indicated at base of bars. Scale bars = 4µm unless indicated otherwise.

**Supplementary Table 1:**
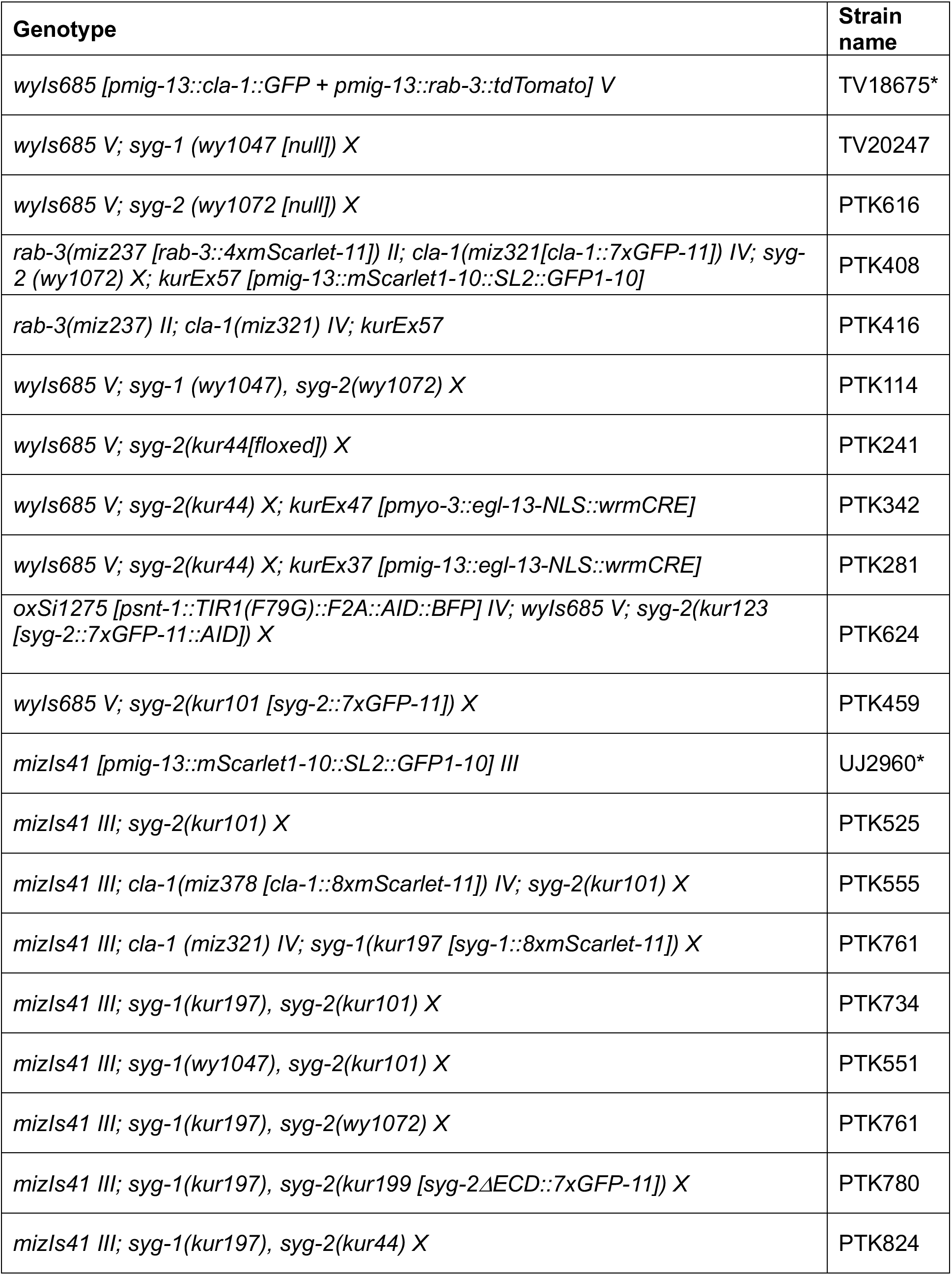
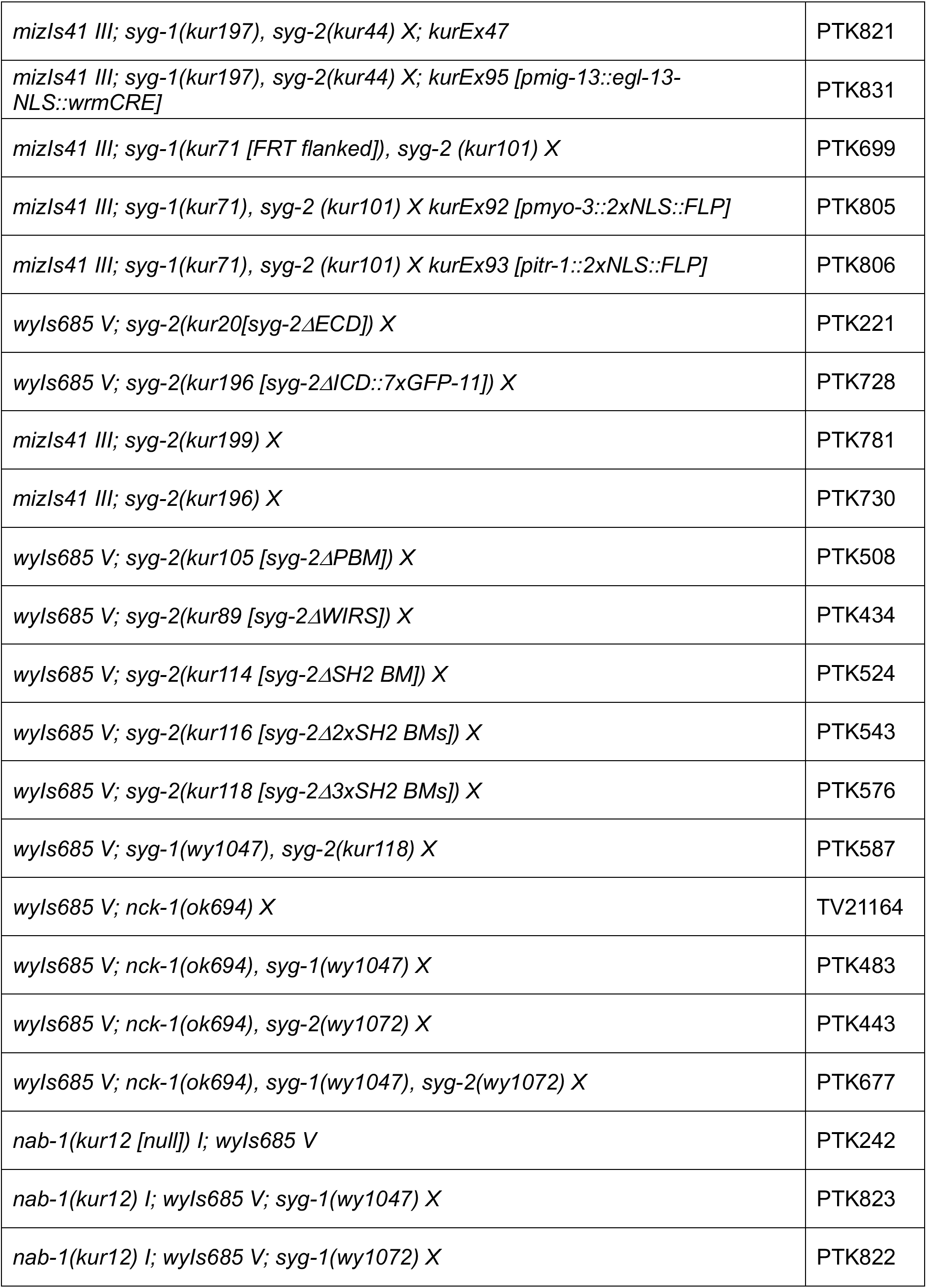
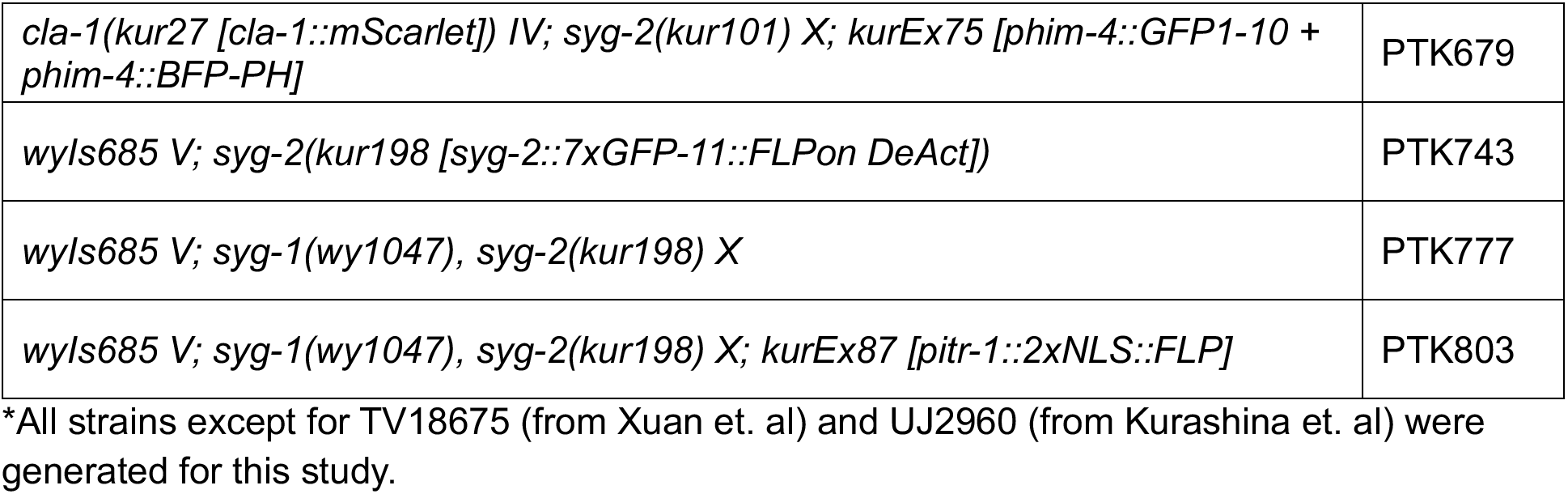
*C. elegans* Strains.

| **Genotype** | **Strain name** |
| --- | --- |
| *wyls685 [pmig-13::cla-1::GFP + pmig-13::rab-3::tdTomato] V* | TV18675* |
| *wyls685 V; syg-1 (wy1047 [null]) X* | TV20247 |
| *wyls685 V; syg-2 (wy1072 [null]) X* | PTK616 |
| *rab-3(miz237 [rab-3::4xmScarlet-11]) II; cla-1(miz321[cla-1::7xGFP-11]) IV; syg-2 (wy1072) X; kurEx57 [pmig-13::mScarlet1-10::SL2::GFP1-10]* | PTK408 |
| *rab-3(miz237) II; cla-1(miz321) IV; kurEx57* | PTK416 |
| *wyls685 V; syg-1 (wy1047), syg-2(wy1072) X* | PTK114 |
| *wyls685 V; syg-2(kur44[floxed]) X* | PTK241 |
| *wyls685 V; syg-2(kur44) X; kurEx47 [pmyo-3::egl-13-NLS::wrmCRE]* | PTK342 |
| *wyls685 V; syg-2(kur44) X; kurEx37 [pmig-13::egl-13-NLS::wrmCRE]* | PTK281 |
| *oxSi1275 [psnt-1::TIR1(F79G)::F2A::AID::BFP] IV; wyls685 V; syg-2(kur123 [syg-2::7xGFP-11::AID]) X* | PTK624 |
| *wyls685 V; syg-2(kur101 [syg-2::7xGFP-11]) X* | PTK459 |
| *mizls41 [pmig-13::mScarlet1-10::SL2::GFP1-10] III* | UJ2960* |
| *mizls41 III; syg-2(kur101) X* | PTK525 |
| *mizls41 III; cla-1(miz378 [cla-1::8xmScarlet-11]) IV; syg-2(kur101) X* | PTK555 |
| *mizls41 III; cla-1 (miz321) IV; syg-1(kur197 [syg-1::8xmScarlet-11]) X* | PTK761 |
| *mizls41 III; syg-1(kur197), syg-2(kur101) X* | PTK734 |
| *mizls41 III; syg-1(wy1047), syg-2(kur101) X* | PTK551 |
| *mizls41 III; syg-1(kur197), syg-2(wy1072) X* | PTK761 |
| *mizls41 III; syg-1(kur197), syg-2(kur199 [syg-2ΔECD::7xGFP-11]) X* | PTK780 |
| *mizls41 III; syg-1(kur197), syg-2(kur44) X* | PTK824 |
| *mizls41 III; syg-1(kur197), syg-2(kur44) X; kurEx47* | PTK821 |
| *mizls41 III; syg-1(kur197), syg-2(kur44) X; kurEx95 [pmig-13::egl-13-NLS::wrmCRE]* | PTK831 |
| *mizls41 III; syg-1(kur71 [FRT flanked]), syg-2 (kur101) X* | PTK699 |
| *mizls41 III; syg-1(kur71), syg-2 (kur101) X kurEx92 [pmyo-3::2xNLS::FLP]* | PTK805 |
| *mizls41 III; syg-1(kur71), syg-2 (kur101) X kurEx93 [pitr-1::2xNLS::FLP]* | PTK806 |
| *wyls685 V; syg-2(kur20[syg-2ΔECD]) X* | PTK221 |
| *wyls685 V; syg-2(kur196 [syg-2ΔICD::7xGFP-11]) X* | PTK728 |
| *mizls41 III; syg-2(kur199) X* | PTK781 |
| *mizls41 III; syg-2(kur196) X* | PTK730 |
| *wyls685 V; syg-2(kur105 [syg-2ΔPBM]) X* | PTK508 |
| *wyls685 V; syg-2(kur89 [syg-2ΔWIRS]) X* | PTK434 |
| *wyls685 V; syg-2(kur114 [syg-2ΔSH2 BM]) X* | PTK524 |
| *wyls685 V; syg-2(kur116 [syg-2Δ2xSH2 BMs]) X* | PTK543 |
| *wyls685 V; syg-2(kur118 [syg-2Δ3xSH2 BMs]) X* | PTK576 |
| *wyls685 V; syg-1(wy1047), syg-2(kur118) X* | PTK587 |
| *wyls685 V; nck-1(ok694) X* | TV21164 |
| *wyls685 V; nck-1(ok694), syg-1(wy1047) X* | PTK483 |
| *wyls685 V; nck-1(ok694), syg-2(wy1072) X* | PTK443 |
| *wyls685 V; nck-1(ok694), syg-1(wy1047), syg-2(wy1072) X* | PTK677 |
| *nab-1(kur12 [null]) I; wyls685 V* | PTK242 |
| *nab-1(kur12) I; wyls685 V; syg-1(wy1047) X* | PTK823 |
| *nab-1(kur12) I; wyls685 V; syg-1(wy1072) X* | PTK822 |
| *cla-1(kur27 [cla-1::mScarlet]) IV; syg-2(kur101) X; kurEx75 [phim-4::GFP1-10 + phim-4::BFP-PH]* | PTK679 |
| *wyls685 V; syg-2(kur198 [syg-2::7xGFP-11::FLPon DeAct])* | PTK743 |
| *wyls685 V; syg-1(wy1047), syg-2(kur198) X* | PTK777 |
| *wyls685 V; syg-1(wy1047), syg-2(kur198) X; kurEx87 [pitr-1::2xNLS::FLP]* | PTK803 |

\*All strains except for TV18675 (from Xuan et. al) and UJ2960 (from Kurashina et. al) were generated for this study.

